# Aphid presence and abundance, more than variation in leaf terpenoid profiles at the plant and plot-level, drive ant behaviour on the perennial forb *Tanacetum vulgare*

**DOI:** 10.64898/2026.02.24.707654

**Authors:** Elikplim Aku Setordjie, Lina Ojeda-Prieto, Wolfgang W. Weisser, Robin Heinen

## Abstract

1) Differences in specialized metabolites are common both within and between plant species and are often thought to have regulating functions in ecological interactions, including herbivores like aphids. Although ants commonly rely on chemical cues in their behaviour and resource finding, very little is known about whether specialized plant chemistry regulates ant behaviour to foster successful ant-aphid mutualisms on plants and in the surrounding vegetation.
2) Using a chemodiversity experiment containing 84 plots with 6 chemotypes of Tansy (*Tanacetum vulgare* L. Asteraceae) planted in different proportions, i.e., plot-level chemotype richness, we tested the effects of plot-level chemotype richness, plot-level chemodiversity metrics, and individual chemotype presence on black garden ant (*Lasius niger* Linnaeus) nesting, patrolling and recruitment behaviour to plants and plots. Furthermore, we assessed the influence of plant chemotype on the pink tansy aphid (*Metopeurum fuscoviride* H.L.G. Stroyan) presence and abundance, as well as ant occurrence on Tansy plants.
3) We found that Tansy plot-level chemodiversity only minimally affected most of the observed ant behaviour, except for nesting, which was marginally positively impacted by plot-level chemotype richness. Clear effects of individual terpenoid chemotypes were observed on ant visitation rates, as well as on aphid presence and abundance. Strongly significant relationships between the probability of ant occurrence and aphid abundance and occurrence observed in our study suggest that ants and aphids are most strongly guided by the presence and abundance of their mutualist partners, rather than by specialized chemistry alone in the Tansy system.

## Introduction

The mutualistic relationship between many ant species (Hymenoptera, Formicidae) and aphids (Hemiptera, Aphididae) is one of the notable examples of mutualism in nature (Way, 1963; Blanchard et al., 2019). Likened by some to animal husbandry by humans, aphids are ‘farmed and milked’ by ants for their sugar-rich excretions, i.e., honeydew, a by-product of feeding on plant phloem sap, which is comparatively rich in sugars, but low in essential amino acids (Douglas et al., 2006; Shaaban et al., 2020). The honeydew that the aphids deposit on plant surfaces attracts sooty mould, which can negatively impact photosynthesis (Insausti et al., 2015; Tedders & Smith, 1976). By collecting honeydew, ants promote aphid colony hygiene (Benton & Foster, 1992; Ivens & Kronauer, 2022). Furthermore, in return for the sugary honeydew, ants often protect aphid colonies from natural enemies (Banks & Macaulay, 1967; Guo et al., 2023); detect and remove sick and dead aphids from colonies (Ivens & Kronauer, 2022); and enable access to preferred plants (Ivens & Kronauer, 2022). This relationship, although beneficial to both mutualistic partners, is often detrimental to their shared plant hosts, as ant-tended aphids suffer from the absence of ants (Flatt & Weisser, 2000). Aphids are commonly considered one of the most destructive plant pests (Guerrieri & Digilio, 2008; Nalam et al., 2019; Twayana et al., 2022; Y. Xu & Gray, 2020). They are highly prolific and are often able to overcome plant defence mechanisms by feeding on phloem sap, and circumventing other plant tissues, which contributes to their detrimental impacts on plant growth (Dixon, 1992; Guerrieri & Digilio, 2008). How exactly ants are able to efficiently find their aphid partners in a complex environment, is a matter of ongoing study (Sakata et al., 2017).

Mutualist ants may locate their aphid partners in several ways, frequently mediated by semiochemicals; specialized compounds that transmit information between organisms of the same and different species (Abd El-Ghany, 2019; Guo et al., 2023; Nault et al., 1976; Verheggen et al., 2012; T. Xu et al., 2021a). For instance, ants can detect a dominant component of aphid alarm pheromones, (E)-β-farnesene (EβF), reorienting their foraging expeditions towards such locations, as they are a reliable cue of aphid presence (Mondor & Addicott, 2007; Nault et al., 1976; Verheggen et al., 2012). This is demonstrated by Verheggen et al. (2012), who showed that in both a choice test and a four arm olfactometer assay *Lasius niger* L. scouts were attracted to *Aphis fabae* alarm pheromone (EβF), even if it was presented in low concentrations. Other studies have found minimal impact of alarm pheromones on ant behaviour (Joachim et al., 2015). It has also been shown that aphids prefer to establish colonies on plants with previous ant presence, suggesting that aphids may also be able to perceive and respond to trail pheromones left behind by ants when foraging (Xu et al., 2021). Plant-derived semiochemicals can also influence ant and aphid behaviour and can potentially influence aphid-associated interactions, including ant-aphid mutualisms. For example Li et al. (2019) observed that young stage-2 pyrethrum flowers produce compounds that mimic aphid alarm pheromone (EβF), recruiting predators for protection and warding off aphids. Junker et al., (2011) also states that *Lasius niger* ants are repelled by natural floral bouquets, due to the presence of monoterpenoids like linalool. Although both ants and aphids have been empirically shown to respond to (plant) metabolites, the influence of plant specialized chemistry on ant nesting, ant patrolling, ant recruitment and ant relationships with aphids is not yet known.

Plants often exhibit chemical diversity, even within species, particularly in their specialized metabolites (Wetzel & Whitehead, 2020; Eilers, 2021; Moore et al., 2014; Ziaja & Müller, 2025). This diversity influences the communication between plants and their environment. Specialized metabolites are essential for plant survival (Akula & Ravishankar, 2011; Weng et al., 2021); they are involved in plant defence against biotic challenges (pests and pathogens) and abiotic stress (environmental conditions e.g. temperature, ultraviolet radiations); they serve as attractants for beneficials during pollination and seed dispersal; and mediate symbiotic relationships with microbes, among others (Kumar et al., 2025; Wink, 2008). Plants are known to produce an astonishing variation in specialized metabolites, and in recent years, it has become clear that the richness, relative abundance and disparity of specialized metabolites, i.e., *chemodiversity*, may guide ecological processes and interactions (Petrén et al., 2024; Wetzel & Whitehead, 2020).

Among the most diverse and abundant specialized metabolites produced by plants are terpenoids, with several thousand known compounds in existence across the plant kingdom, which can be produced either constitutively or when induced by biotic and abiotic stress (Broun & Somerville, 2001). Specialized metabolites produced constitutively are stored in plant tissues and emitted either by evaporation or when plant tissues are disrupted mechanically. These compounds make up a plant’s chemotype, which may be detected by plant associated organisms like herbivores, natural enemies, among others, resulting in attraction or repulsion (Clancy et al., 2016). Induced specialized metabolites, however, are produced and emitted in direct response to stressors e.g. herbivory (Clancy et al., 2016; Dicke & Van Loon, 2000; Pare & Tumlinson, 1997). Terpenoids are the largest group of specialized metabolites produced by plants, many of which play a role in plant defence against biotic stress, and depending on the metabolite and their metabolic class, can be stored in tissues or released as volatiles (Kumar et al., 2025; Clancy et al., 2016; Newrzella et al., 2025). Terpenoids have varying effects on herbivore feeding and abundance via different pathways (Kleine & Müller 2013; Jakobs et al. 2018; Li et al., 2023; Neuhaus-Harr *et al*. 2024; 2025). For instance, terpenoids within Lemon (*Citrus limon*) fruit peel aqueous extract have been observed to be highly toxic to rose aphid (*Macrosiphum roseiformis)* both in lab and field assays (Gupta et al., 2017). In lab-based pairwise choice assays, Neuhaus-Harr et al. (2024) also found that Tansy aphid (*Macrosiphoniella tanacetaria*) had a clear preference and attraction to Tansy (*Tanacetum vulgare*) leaflets with terpenoid profiles that were dominated by a mixture of α-thujone and β-thujone or by trans-chrysanthenyl acetate, whilst avoiding leaflets with terpenoid profiles dominated by α-pinene and sabinene, although these chemotypes did not correspond to the chemotypes on which the aphid colonies performed best (Neuhaus-Harr et al., 2025).

The ecological consequences of specialized chemical diversity can be studied at different scales, from the plant tissue scale to that of entire plant individuals, and up to the variation in individuals at the community or population level (Calixto et al., 2025; Clancy et al., 2016; Ojeda-Prieto, Medina-van Berkum, et al., 2025). However, to date only few studies have focused on chemodiversity at these higher levels of biological organization, and the consequences it may have for interactions at larger spatial scales (Hanusch et al., 2026). Plant chemodiversity can be manipulated at plot-level by growing plants with different individual chemical profiles within the same plot, using different combinations of plant chemotypes to represent different levels of chemical richness (Ojeda-Prieto, Moreno, et al., 2025; Ojeda-Prieto, Medina-van Berkum, et al., 2025). Plot-level manipulations elicit complex responses in plant neighbours and associated organisms (Bustos-Segura et al. 2017; Sasidharan et al., 2024; Ziaja & Müller, 2023). However, the impacts of population-level chemodiversity on mutualistic interactions between ants and aphids have received minimal attention to date (but see Rahimova et al., (2024) for a study linking terpenoid chemistry to interactions at large geographic scales).

*Tanacetum vulgare* L., commonly known as Tansy, is an herbaceous plant of the Asteraceae family that exhibits high intraspecific diversity in mono- and sesquiterpenoids (Kleine & Müller, 2013; Rahimova et al., 2024). This diversity influences its interactions with associated herbivores (Clancy et al., 2020; Neuhaus-Harr et al., 2024; Ojeda-Prieto, Moreno, et al., 2025; Senft et al., 2019). Tansy plant individuals can be grouped into different chemotypes based on hierarchical clustering analysis of their leaf terpenoid blends (Ojeda-Prieto, Medina-van Berkum, et al., 2025). Several studies have investigated the effects of chemodiversity and chemotype richness on Tansy associated herbivores at plant and plot-levels. According to Ojeda-Prieto, Moreno, et al. (2025), Tansy chemodiversity and chemotype richness have negative effects on herbivore abundance; a relationship largely driven by the responses of the crimson tansy aphid (*Uroleucon tanaceti*). *Metopeurum fuscoviride,* commonly known as the pink tansy aphid, is another specialist herbivore of the Tansy plant. It is an obligate myrmecophile, predominantly tended to by the black garden ant (*Lasius niger*) (Stadler et al., 2002). As observed by (Ojeda-Prieto, Moreno, et al., 2025), *M. fuscoviride* has variable responses to plant and plot-level chemotype richness; it responds negatively to increasing chemotype richness in some years, but has no response in other years. Given its intimate relationship with *L. nige*r, it is plausible that plant and plot-level chemodiversity and chemotype richness might influence *L. niger* behaviour and, in turn, their mutualistic aphid partners. However, little is known about these effects, hence, this study aims to understand the relationships between plant- and plot-level chemodiversity on the nesting, patrolling and recruitment behaviour of mutualistic ant *Lasius niger* and *M. fuscoviride* aphid presence and abundance.

With *Tanacetum vulgare* as our model plant, we designed a field experiment with plots containing six plants of varying numbers of chemotypes, i.e., 1, 2, 3 and 6 chemotypes per plot. In the 2024 summer season, we observed the number of nests within a meter of plots and the number of ants patrolling within plots in the absence of bait. We also investigated recruitment of ants to sugar baits in the plots. Finally, we examined the occurrence *L. niger* individuals and the occurrence and abundance of *M. fuscoviride* individuals on each Tansy plant in each plot.

With these data we aimed to test the following research questions:

1) Does plot-level chemotype richness affect ant nesting, patrolling and recruitment behaviour?
2) Do plot-level chemodiversity metrics (chemical richness, chemical abundance, chemical Shannon diversity, chemical evenness) affect ant nesting, patrolling and recruitment behaviour?
3) Does the presence of specific chemotypes in a plot affect ant nesting, patrolling and recruitment behaviour?
4) Do specific chemotypes differ in their attractiveness to ants and aphids, and are these linked?

## Materials and Methods

### Experimental field

A chemodiversity field experiment has been running within the grounds of the Jena Experiment, in Jena, Germany, since 2021 (described in detail in Ojeda-Prieto, Medina-van Berkum, et al., (2025), see also Fig. 1). Briefly, 84 plots were planted with different combinations of six selected Tansy (*Tanacetum vulgare*) chemotypes differing in their leaf terpenoid profiles. Chemotypes were named by the characteristics of their profiles, in terms of dominant compounds and respective concentrations (i.e., “Athu-Bthu”, “Bthu-low”, “Bthu-high”, “Mixed-high”, “Mixed-low” and “Chrys-acet”) (Neuhaus-Harr et al., 2024; Ojeda-Prieto, Medina-van Berkum, et al., 2025).

**Figure 1:**
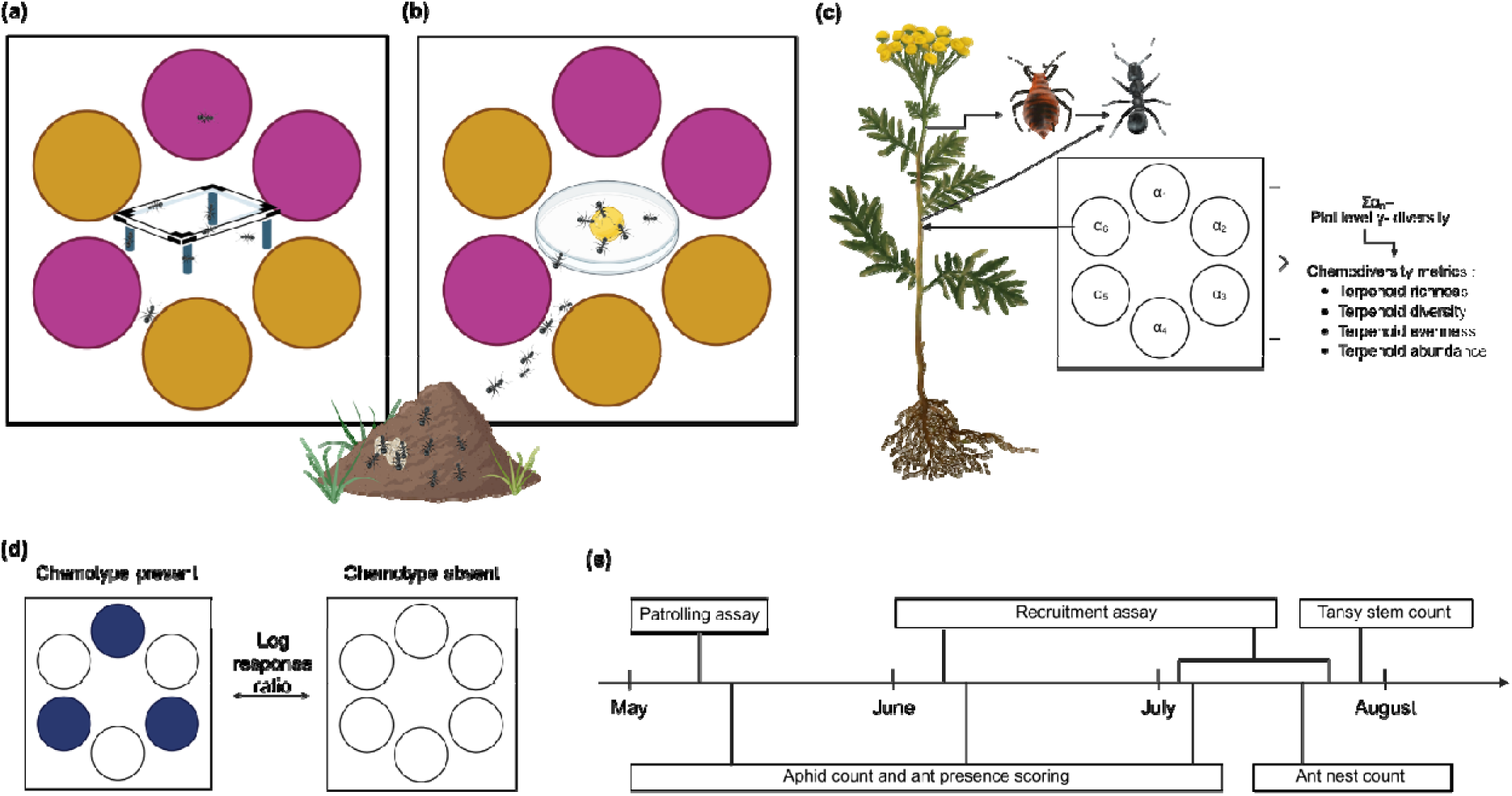
Experimental design and timeline of the field experiment (a) illustration of ant patrolling assay where a square quadrat is placed in the centre of a plot; (b) illustration of recruitment assay where honey on a petri dish cover is placed at the centre of a plot and an ant nest found at the edge of plot; (c) illustration of plot chemodiversity metrics derived from Tansy leaf samples and its effects on aphids and ants on plants, where α represents the terpenoid diversity of each individual plant within plot and plot γ diversity represents the sum of terpenoid diversity of all plants within plot; (d) illustration of log response ratio where ant behaviour when a chemotype is present within plot is compared to when a chemotype is absent; (e) timeline of the field survey in 2024. NB: circles represent Tansy plants within plots; different coloured circles represent different Tansy chemotypes. Created with Biorender® Agreement number: FQ29ECFJ5L.

Plots were planted with six plants, with varying plot-level chemotype richness levels of 1, 2, 3, or 6 chemotypes planted per plot. Each level was replicated 12, 30, 30 and 12 times respectively. Plots of each level were equally distributed and randomly placed within six spatial blocks containing 14 plots. Each plot was 1 x 1 m, divided by 70cm wide paths that were mown regularly in summer. For each chemotype, three biologically unique daughter genotypes were cloned via cuttings, these 18 daughters are present 27-31 times in different combinations across the different plots in the field.

### Ant nest counts

To investigate the distribution of ant nests across the chemodiversity experiment, the *number of nests within one metre of the plot* (visible ant nest openings) measured from the centre of plots were assessed in July 2024, as a proxy for local ant nest density. We looked for nests by lightly and superficially scratching the soil surface in and around plots to stimulate ants, followed by scanning for activity, and by checking activity in mounds. An opening was only recorded if either eggs and larvae were observed, or when more than ten mature ants emerged after the soil was scratched.

### Ant patrolling behaviour

To investigate ant patrolling behaviour in plots across the chemodiversity experiment, the *number of ants patrolling* within plots was recorded in the absence of external baits once, on the 16^th^ of May 2024, between 9:30am and 4:30pm at a mean atmospheric temperature of 19.7°C, when *T. vulgare* plants were in the early growth stage. The observation was done using a square quadrat of area 32.5 x 32.5 cm, which was raised above the soil at 2.5 cm to prevent interference with ant patrolling. The quadrat was placed in the centre of each plot, and observed for 5 minutes, during which the number of ants walking through the quadrat were recorded. Ants were counted once they entered the quadrat, i.e., ants that left the quadrat and entered again were considered additional patrolling activity within the quadrat (Fig. 1a).

### Ant recruitment-to-bait behaviour

To investigate differences in ant recruitment across the chemodiversity experiment, sugar baits were used to observe differences in ant recruitment to the baits over time. To this end, Petri dish covers of 14 cm diameter were placed, upside down, in the centre of each plot. After placement of the dishes, 2 ml of honey (Langnese Flotte Biene, Bargteheide, Germany) was placed at the centre of each, using plastic syringes. After the honey placement, ant numbers were counted five times at 10-minute intervals. All recruitment counts were done block by block. The approximate time used for placing petri dishes and adding honey in all 14 plots within a block was two minutes. From these ant count data, we assessed the *time to first ant recruitment*, i.e., a plot with their first ants in the first count was given a recruitment time of 10 minutes. Plots with no ants at the end of the five counts were given a time point of 60, indicating that they had a recruitment time of at least 60 minutes. The *total number of ants* on the honey baits was determined by summing up the number of ants on a bait across all five counts (Fig. 1b).

The ant recruitment assays were conducted three times, on 07^th^ June 2024 (i.e. early June between 11:00 AM and 5 PM at a mean atmospheric temperature of 21.7°C) and 5^th^ July, 2024 (early July between 9:00 AM to 2:30 PM at a mean atmospheric temperature of 19.4°C), the counts were replicated twice on both days by two different observers, each observing three blocks per replicate round, and in late July with the first replicate round on the 24^th^ (8:30 AM – 1:30 PM, mean atmospheric temperature = 20.2°C) and the second on 25^th^ of July 2024 (8:30 AM – 2:00 PM, mean atmospheric temperature = 22.6°C), by the same observer. Two ant species were recorded on the honey baits: *Lasius niger* and *Formica rufibarbis*. Although statistical analyses are conducted on data from both species, as *L. niger* was the most common visitor (89.6 % of counts), further discussions are made mentioning only this species.

### Temperature

As ants in temperate climates tend to be thermophilic, differences in behaviour are to be expected, depending on time of the day and the weather conditions on the respective sampling days. To investigate the effects of temperature on ant-aphid dynamics in the chemodiversity experiment at different times of the year and day, temperature was estimated using the atmospheric temperature of the approximate time of day that ants were counted in each plot during the ant recruitment assay. This data, taken at 30-minute intervals, was supplied by the weather station at <u>The Jena Experiment</u>.

### Aphid counting

The number of pink tansy aphid (*Metopeurum fuscoviride)* individuals on each plant was recorded three times, in the months of May (16^th^ - 17^th^), June (6^th^) and July (4^th^) of 2024. The presence of ants on each plant in these months was also recorded at the same time. For plants with smaller aphid colony numbers, the number of aphids on each stem was counted, when aphid colonies numbers were higher (>100), aphid numbers were estimated by counting aphids on 1 cm of stem on three different stems and multiplying by the length of the stem that was colonized, and multiplied by plant stem number; all aphid counts above 500 were recorded as 500 individuals.

### Plant stem numbers

To account for the effect of Tansy plant density on ant-aphid dynamics across the chemodiversity experiment, the number of stems per plant was recorded once in July of 2024 and then averaged by the number of plants present in the plot (i.e., 6) to be used as a covariate.

### Statistical analyses

Data was analysed using R version 4.5.0 (R Core Team, 2025). The (fixed) effect of plot-level chemotype richness on the number of nests within a meter of plot, ant patrolling, the time to first ant recruitment, and the total number of ants on sugar bait was assessed with linear mixed effects models using the *lme4* package (Bates et al., 2015).

For the number of nests within a meter of plot, which was recorded once, only block was included as a random effect to account for the spatial structure of the field experimental design:

> **Model 1:** lmer (Number of nests within a meter of plot ∼ Average number of stems within plot + Plot-level chemotype richness + (1|Block))

A similar model structure was used for the number of ants patrolling within plots which was also recorded once:

> **Model 2:** lmer (Number of ants patrolling ∼ Number of nests within a meter of plot + Average number of stems within plot + Plot-level chemotype richness + (1|Block))

For time to first ant recruitment and for ant abundance, which were repeated in different months and repeat rounds, we added plot nested within block as random effects to account for the spatial and temporal structure of the sampling design.

> **Model 3:** glmer (Time to first recruitment ∼Number of nests within a meter of plot + Temperature + Average number of stems within plot + Month * Plot-level chemotype richness+(1|Block/Plot), family = poisson)
>
> **Model 4:** lmer (log (Total number of ants on honey baits +1) ∼ Number of nests within a meter of the plot + Temperature + Average number of stems within plot + Month * Plot-level chemotype richness + (1|Block/Plot)

For each model, residual distributions were checked using diagnostic plots, and data were transformed, where necessary, to approach normality and homoscedasticity of the residuals.

Plot-level chemodiversity metrics, i.e. chemical richness, Hill evenness (q = 1), abundance and Shannon diversity, were calculated based on the individual leaf terpenoid profiles of every plant individual present in a plot and analysed with the ‘chemodiv’ package (Petrén et al., 2023). Note that the chemical analyses were done using leaf material from the original chemotype lineages, which were clonally propagated for the field establishment in 2021 (Ojeda-Prieto, Medina-van Berkum, et al., 2025). Since then, these lineages have been chemotyped multiple times and have been shown to be conserved in their chemical composition, even between experimental years (Newrzella et al., 2025). We therefore used the original profiles to forecast plot-level chemical diversity proxies, which can provide insights into the relationships between plot-level chemodiversity and plot-level ecological functioning within this field experiment. The effect of *terpenoid richness*, *terpenoid Shannon diversity*, *terpenoid Hill evenness*, and *relative terpenoid abundance* on ant nesting, ant patrolling, time to first ant recruitment and total number of ants on bait was assessed using linear mixed models with the *lme4* package with a random effect structure as described above. To make predictor variables more comparable, predictor variables in Model 6 – 8 were standardized to a mean of 0 and a standard deviation of 1 using the scale () function in R.

> **Model 5:** lmer (Number of nests within a meter of the plot ∼ Average number of stems within plot + Terpenoid abundance +Terpenoid Hill Evenness + Terpenoid Richness + (1|Block))
>
> **Model 6:** lmer (Number of ants patrolling ∼ Number of nests within a meter of the plot + Average number of stems within plot + Terpenoid abundance + Terpenoid Hill Evenness +Terpenoid richness + (1|Block))
>
> **Model 7:** lmer (log (Total number of ants on honey bait +1) ∼ scale (Number of nests within a meter of plot) + scale (Temperature)+ scale (Average number of stems within plot) + scale (Terpenoid abundance) *Month + scale (Terpenoid Hill Evenness) + scale (Terpenoid richness) + (1|Block/Plot))
>
> **Model 8:** glmer (Time to first recruitment ∼ scale (Number of nests within a meter of plot) + scale (Temperature) + scale (Average number of stems within plot) + scale (Terpenoid abundance) *Month + scale (Terpenoid Hill Evenness) + scale (Terpenoid richness) + (1|Block/Plot), family = poisson)

We tested for multicollinearity between explanatory variables using variance inflation factors. Shannon diversity was excluded from the model due to strong correlation with Hill evenness.

The effect of the presence and absence of individual Tansy chemotypes on ant nesting, ant patrolling, time to first ant recruitment, and total number of ants on bait was analysed with linear mixed effects models using the *lme4* package with a random effects structure as described above:

> **Model 9:** lmer (Number of nests within a meter of plot ∼ Average number of stems within plot + Chemotype + (1|Block))
>
> **Model 10:** lmer (Number of ants patrolling ∼ Number of nests within a meter of plot + Average number of stems within plot+ Chemotype + (1|Block))

Because several variables were assessed more than once over time, we also included plot as a random factor for these variables:

> **Model 11:** lmer (log (Total number of ants on honey bait + 1) ∼ Number of nests within a meter of plot + Average number of stems within plot + Temperature + Chemotype *Month + (1|Block/Plot))
>
> **Model 12:** glmer (Time to first recruitment ∼ Number of nests within a meter of plot + Average number of stems within plot + Temperature + Chemotype *Month + (1|Block/Plot), family = poisson)

Where Chemotype was 1 when present within plot and 0 when absent within plot.

To investigate the effect of chemotype on aphid and ant incidence on Tansy plants, ant occurrence, aphid abundance and aphid occurrence were averaged across all Tansy clonal daughters (i.e., technical replicates) present in different plot combinations within our field. This means that there were six chemotypes with three true biological replicates each. One-way ANOVA was used to test the effect of chemotype on average ant occurrence, average aphid occurrence and average aphid abundance. When significant, Tukey’s HSD post hoc test was used to identify differences between chemotypes using the *emmeans* and *multcomp* packages (Lenth & Piaskowski, 2017; Hothorn et al., 2008). After which the relationship between the various means were determined using simple linear regressions (R Core Team, 2025).

All R scripts, including model specifications are provided in the supplementary information to this manuscript. Data was organized using *dplyr* (Wickham et al., 2023). Graphics were created using academic-licensed Biorender®, *ggplot2* and *ggpubr* (Wickham, 2016; Kassambara, 2025). All P-values were calculated using the Anova function in the *car* package (Fox et al., 2019).

## Results

Over the course of sampling, we recorded a total of 5864 ants in our field plots. 1617 were recorded patrolling within square quadrats in the absence of bait in May, whilst 4247 ants, i.e. 3807 *Lasius niger* and 439 *Formica rufibarbis*, were recorded on sugar baits over three sampling time points in early June, early July and late July. For aphids, approximately 11795 *Metopeurum fuscoviride* individuals were recorded on the Tansy plants in our 84 plots in May, early June and early July.

### Effect of plot-level chemotype richness on ant behaviour

The number of ant nests observed within one meter of plots increased with increasing average number of stems within plots (X² = 4.6, p = 0.032, Table S1). Plot-level chemotype richness only had a marginally positive effect on the number of nests within a meter of the plot (X² = 3.6, p = 0.058, Fig. 2a).

**Figure 2:**
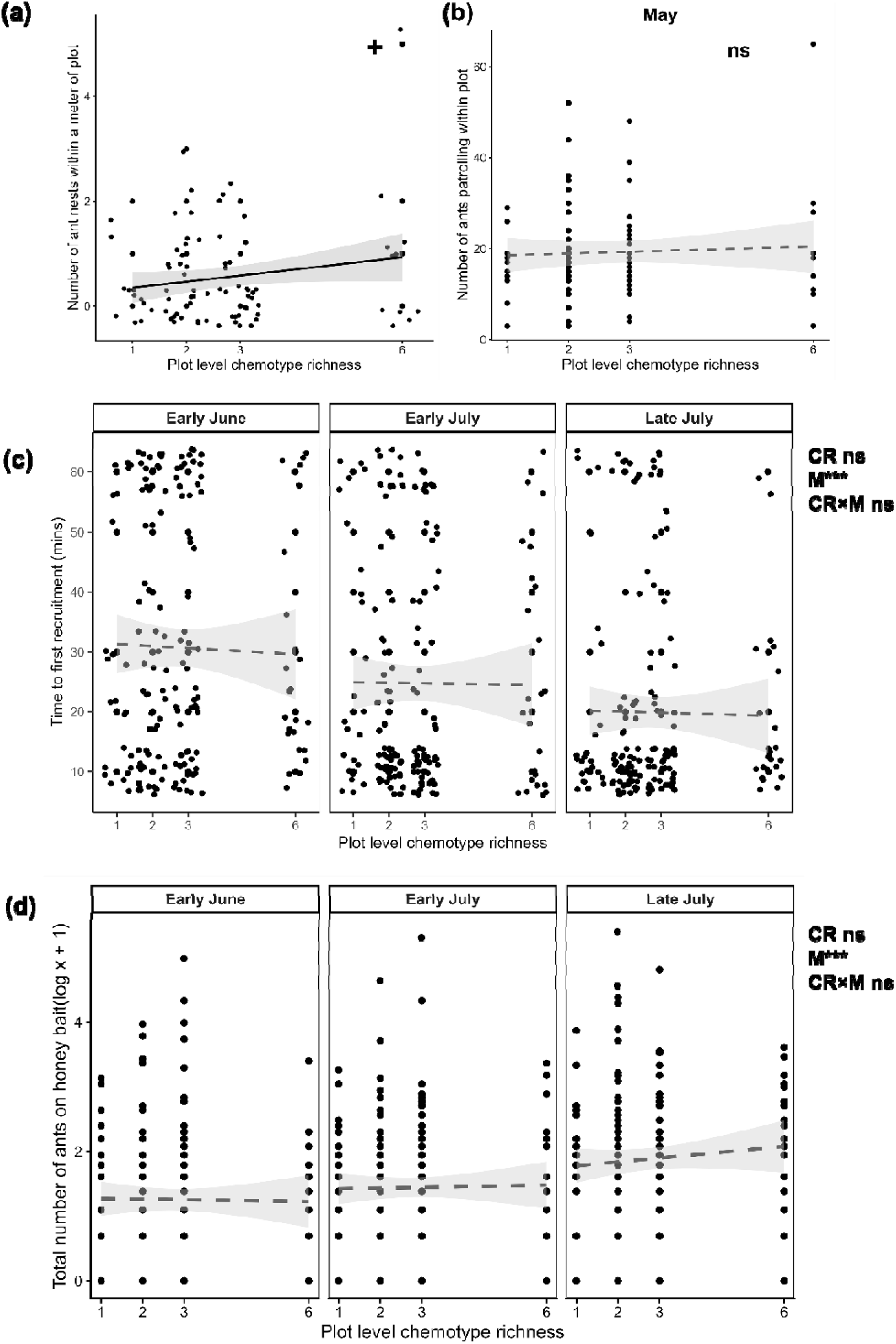
The effect of *Tanacetum vulgare* plot-level chemotype richness (CR), month of observation (M) and their interaction on (a) number of nests within a meter of plot, (b) number of ants patrolling within plots in the absence of bait, (c) time to first ant recruitment on baits and (d) total number of ants on baits. Dashed lines represent non-significant results; solid lines represent (marginally) significant results. Significance levels are presented as: ns = not significant, + = 0.10 > *P* > 0.05, * P<0.05, ** P<0.01, and *** P<0.001 on the top right corner of figure. Data points (dots) are jittered in (c) for better visualisation.

The number of ants patrolling within plots in the absence of bait was significantly higher when more nests were observed within a meter of the plot (X² = 7.2, p = 0.007, Table S1). However, average number of stems within plots (X² = 0.8, p = 0.365, Table S1) and plot-level chemotype richness did not significantly affect the number of ants patrolling within plots in the absence of bait (X² = 0.0, p = 0.962, Fig. 2b).

Ant recruitment was influenced by atmospheric temperature (X² = 31.3, p = <0.001, Fig. S2). Specifically, recruitment times decreased from the first to last sampling times. However, within early June and early July higher temperature increased the time to first recruitment to baits whilst in late July higher temperatures lead to faster ant recruitment. Average number of stems in plot (X² = 0.7, p = 0.393, Table 1), number of nests within a meter of plot (X² = 1.1; p = 0.293; Table S1) and plot-level chemotype richness (X² = 0.0, p = 0.912, Fig. 2c) did not impact time to first recruitment on baits. The interaction between month and plot-level chemotype richness was also not significant (X² = 0.3, p = 0.849, Table S1).

Temperature (X² = 0.0, p = 0.914, Table 1), average number of stems in plot(X² = 1.4, p = 0.239, Table S1), number of nests within a meter of plot (X² = 0.0; p = 0.831; Table S1) and plot-level chemotype richness (X² = 0.2, p = 0.623, Fig. 2d) did not significantly affect the total number of ants found on baits across all observed time points. Neither was there a significant interaction between month and plot-level chemotype richness (X² = 1.1, p = 0.583, Table S1).

### Effects of plot-level chemodiversity metrics on ant behaviour

Nest numbers within a meter of plots were positively affected by the average number of stems within plots (X² = 4.5, 0.033, Table S2). Plot-level terpenoid richness (X² = 0.5, p = 0.482, Table S2), abundance (X² = 0.9, p = 0.350, Fig. 3a) and Hill evenness (X² = 0.1, p = 0.713, Table 2), however, had no effect on nest numbers within a meter of plot.

**Figure 3:**
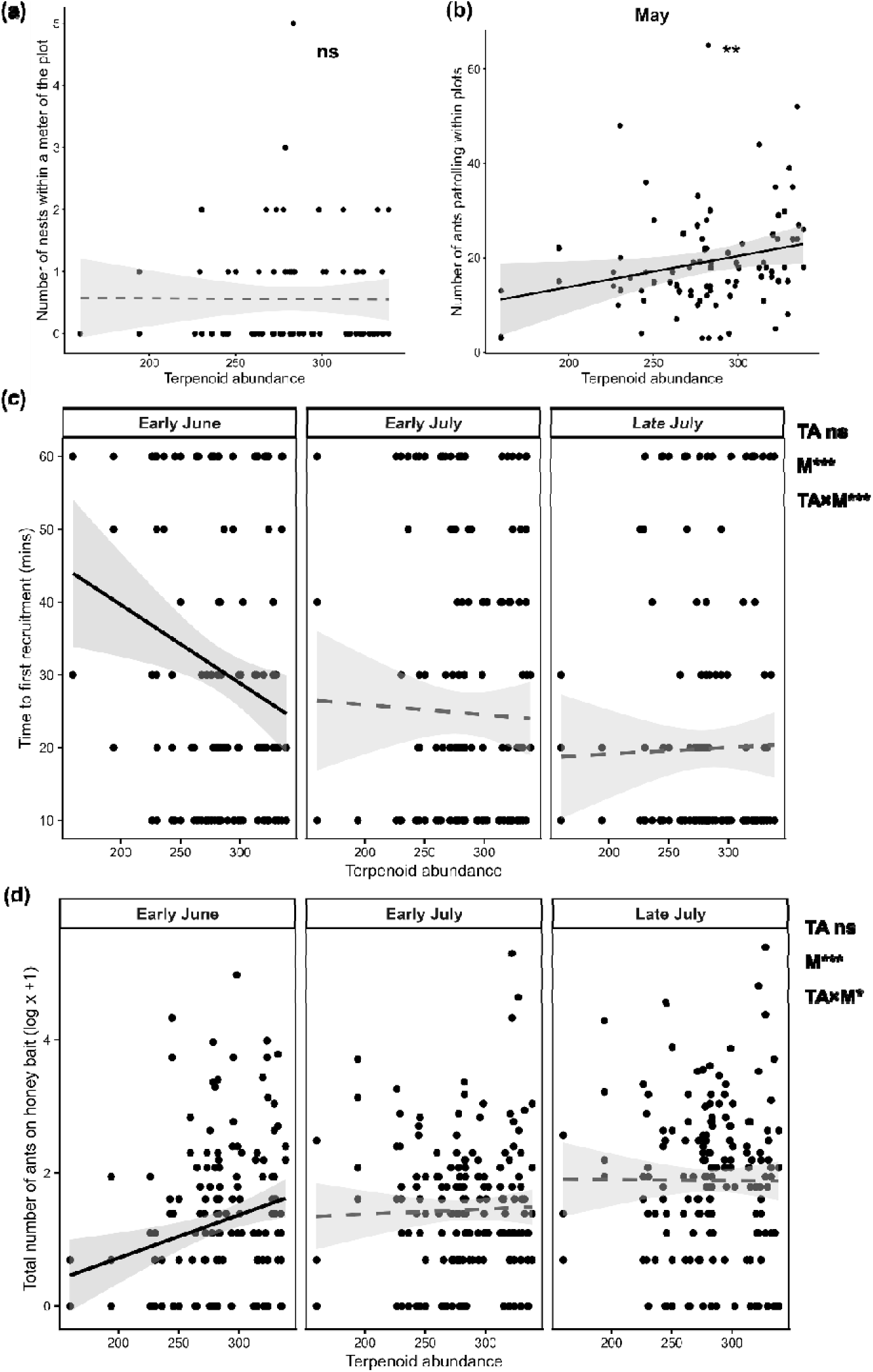
The effect of Terpenoid abundance (TA) and month of observation (M), and their interaction on (a) number of nests within a meter of plot, (b) number of ants patrolling within plots in the absence of bait, (c) time to first ant recruitment on baits and (d) total number of ants on honey baits. Dashed lines represent non-significant results; solid lines represent (marginally) significant results. Significance levels are presented as: ns = not significant, + = 0.10 > *P* > 0.05, * P<0.05, ** P<0.01, and *** P<0.001 on the top right corner of figure.

The average number of stems within plots (X² = 0.5, p = 0.465) did not affect the number of ants patrolling within plots but plots having higher number of nests within a meter also had higher ant patrolling (X² =6.0, p = 0.014). Plot-level terpenoid richness (X² = 0.4, p = 0.510, Table S2) and Hill evenness (X² = 0.0, p = 0.983, Table 2) did not influence the number of ants patrolling within plots in the absence of bait, except for relative terpenoid abundance (X² = 10.4, p = 0.001, Fig. 3b), which had a positive relationship with the number of ants patrolling within plots.

Slower ant recruitment to baits was observed at higher atmospheric temperatures in early June and early July whilst higher temperature in late July resulted in faster recruitment (X² = 33.3, p = <0.001, Fig. S2). Average stem numbers within plot (X² = 0.0, p = 0.828) and the number of nests found around plots (X² = 0.9, p = 0.349) had no impact on the rate of ant recruitment to baits. Time to first recruitment was not influenced by plot-level terpenoid richness, terpenoid Hill evenness and terpenoid abundance (Figure 3c and 3d, Table S2). A significant interaction was observed, however, between terpenoid abundance and month (X² = 50.9, p = <0.0001, Fig. 3c) on time to first recruitment, where higher terpenoid abundance in plots resulted in faster recruitment to baits in early June, an effect that gradually faded in subsequent months. We also observed a significant interaction between terpenoid Hill evenness and month (X² = 10.9, p = 0.004, Table 2, Fig. S3) on time to first recruitment, higher terpenoid Hill evenness increased time to first recruitment in early June and early July but no clear effect on time to first recruitment in late July.

The total number of ants recorded on baits was not affected by temperature (X² = 0.0, p = 0.967), the average number of stems within the plot (X² = 0.4, p = 0.519) and the number of nests within a meter of the plot (X² = 0.1, p = 0.706). Plot terpenoid richness (X² = 2.1, p = 0.143), Hill evenness (X² = 1.6, p = 0.212) and Abundance (X² = 0.1, p = 0.704) also had no effects on the total number of ants on baits but a significant interaction was observed between terpenoid abundance and month (X² = 7.3, p = 0.026). In early June, higher terpenoid abundance in plots positively influenced the total number of ants on baits, but there was no difference in early July and late July (Fig. 3d, Table S2).

### Effects of specific (individual) chemotypes on ant behaviour

The average number of stems with plots had significant effects on the number of nests found within a meter of plot in the presence of all chemotypes (Table S3). In the presence of the Chrys-acet (X² = 4.0, p = 0.045, LRR = 0.58, Fig. 4a) and Bthu-low (X² = 2.7, p = 0.099, LRR = 0.42, Fig. 4a) chemotypes, the mean number of nests found within a meter plot increased by 78.6 %. and 52.2 % respectively, compared to plots where those chemotypes were absent. All other plant chemotypes did not significantly affect nest numbers around plots (Table 3).

**Figure 4:**
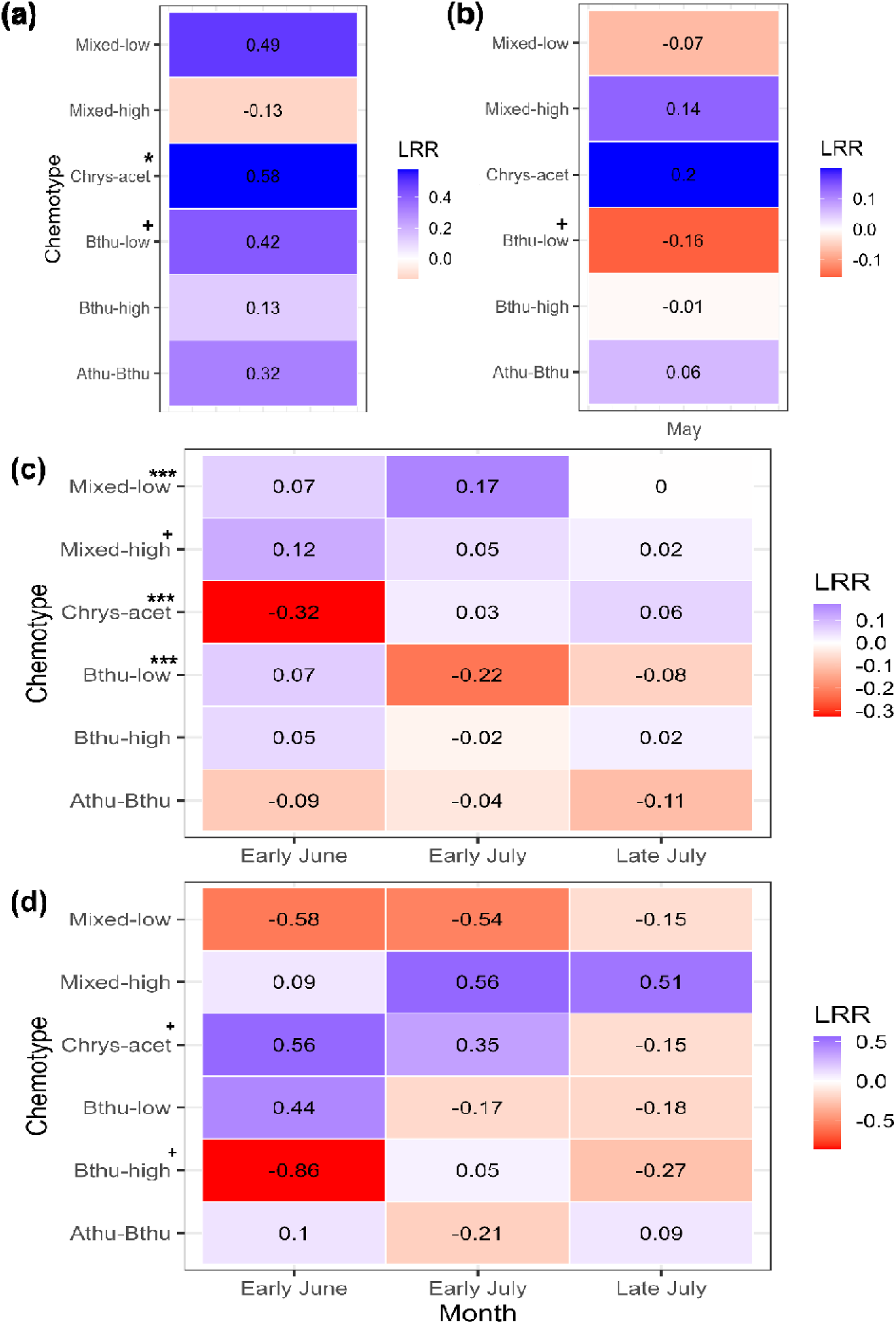
The effect of the presence of individual plant chemotypes on log response ratios (LRR) of (a) number of nests within a meter of the plot, (b) number of ants patrolling within plots in the absence of bait, (c) time to first ant recruitment on baits and (d) total number of ants on honey baits. Significance levels are presented as: + = 0.10 > P > 0.05, * P<0.05, ** P<0.01, and *** P<0.001. Significant interactions between chemotype and month indicated on top of chemotype names in c and d. NB: Log response ratio (LRR) compares the response of ant behaviour when a chemotype is present to when a chemotype is absent (i.e. ln (mean response in presence of chemotype /mean response in absence of chemotype). Colour gradient between blue (positive response to chemotype presence), white (no effect) and red (negative response to chemotype presence).

The number of ants patrolling within plots was significantly higher in plots having more nests within a meter radius regardless of chemotype presence or absence but was unaffected by the average number of stems within plots (Table S3). Nevertheless, the number of ants patrolling within plots was marginally negatively impacted by the presence of the Bthu-low chemotype (X² = 2.7, p = 0.099, LRR = −0.16, Fig. 4a) compared to plots without the chemotype, the presence of all other plant chemotypes did not significantly impact ant patrolling within plots (Fig. 4b, Table S3).

Atmospheric temperature influenced ant recruitment within plots in the presence and absence of all chemotypes whilst average number of stems and the number of nests within a meter of plot had no impact on time to first recruitment (Table S3). The presence of individual chemotypes within plots also had no significant main effect on the time to first ant recruitment on baits (Table 3). Although, significant interactions were observed between month and the presence of specific chemotypes: Chrys-acet (X² = 98.7, p = <0.001), Bthu-low (X² = 47.5, p = < 0.001), Mixed-low (X² = 14.7, p = <0.001) and Mixed-high (X² = 5.1, p = 0.074). For the plots in which the Chrys-acet chemotype was present, time to first recruitment increased over time, compared to plots in which the chemotype was absent (Fig.4c). Bthu-low presence in plots initially led to longer time to first recruitment in early June, this changed in subsequent months where the presence of the chemotype led to faster recruitment with the fastest recruitment time observed in early July, when compared to plots without the chemotype (Fig 4c). Time to first ant recruitment though varying from month to month in the presence of the Mixed-low and Mixed-high chemotypes was relatively longer compared to other plots that did not contain these chemotypes (Fig 4c).

Temperature, average number of stems within plot and the number of nests within a meter of plot had no effect on the total number of ants found on baits in the presence and absence of all chemotypes (Table S3). Similarly, the presence of individual plant chemotypes had no significant main effect on the total number of ants found on baits (Fig. 4d, Table S3). However, significant interactions were observed between month and two chemotypes, i.e. Chrys-acet (X² = 5.8, p = 0.054) and Bthu-high (X² = 4.9, p = 0.088). The presence of the Chrys-acet chemotype positively influenced the number of ants on baits in early June, this positive effect however decreased as the months progressed, by late July the presence of this chemotype negatively affected ant numbers on baits (Fig. 4c). An opposite pattern was observed with the Bthu-high chemotype which negatively affected ant numbers on baits in early June but had the negative effect reducing in subsequent months (Fig. 4c).

### Attractiveness of specific chemotypes to ants and aphids

Average *Lasius niger* occurrence (F_5,12_ = 3.5, p = 0.037) and average *Metopeurum fuscoviride* abundance were both significantly affected by Tansy chemotype (F_5,12_ = 8.8, p = 0.001). Aphid occurrence, however, was only marginally affected by chemotype (F_5,12_ = 2.9, p = 0.061). Both *M. fuscoviride* and *L. niger* showed a clear preference for the Mixed-high chemotype compared to all other chemotypes, whilst Mixed-low had the lowest incidence of both ants and aphids. For some chemotypes like Chrys-acet, *L. niger* occurrence and *M. fuscoviride* occurrence and abundance differed between replicate daughters. The presence and abundance of *M*. *fuscoviride* on plants had a significantly positive impact on *L. niger* occurrence. *M*. *fuscoviride* occurrence and abundance also had a significantly positive relationship (Fig. 5).

**Figure 5:**
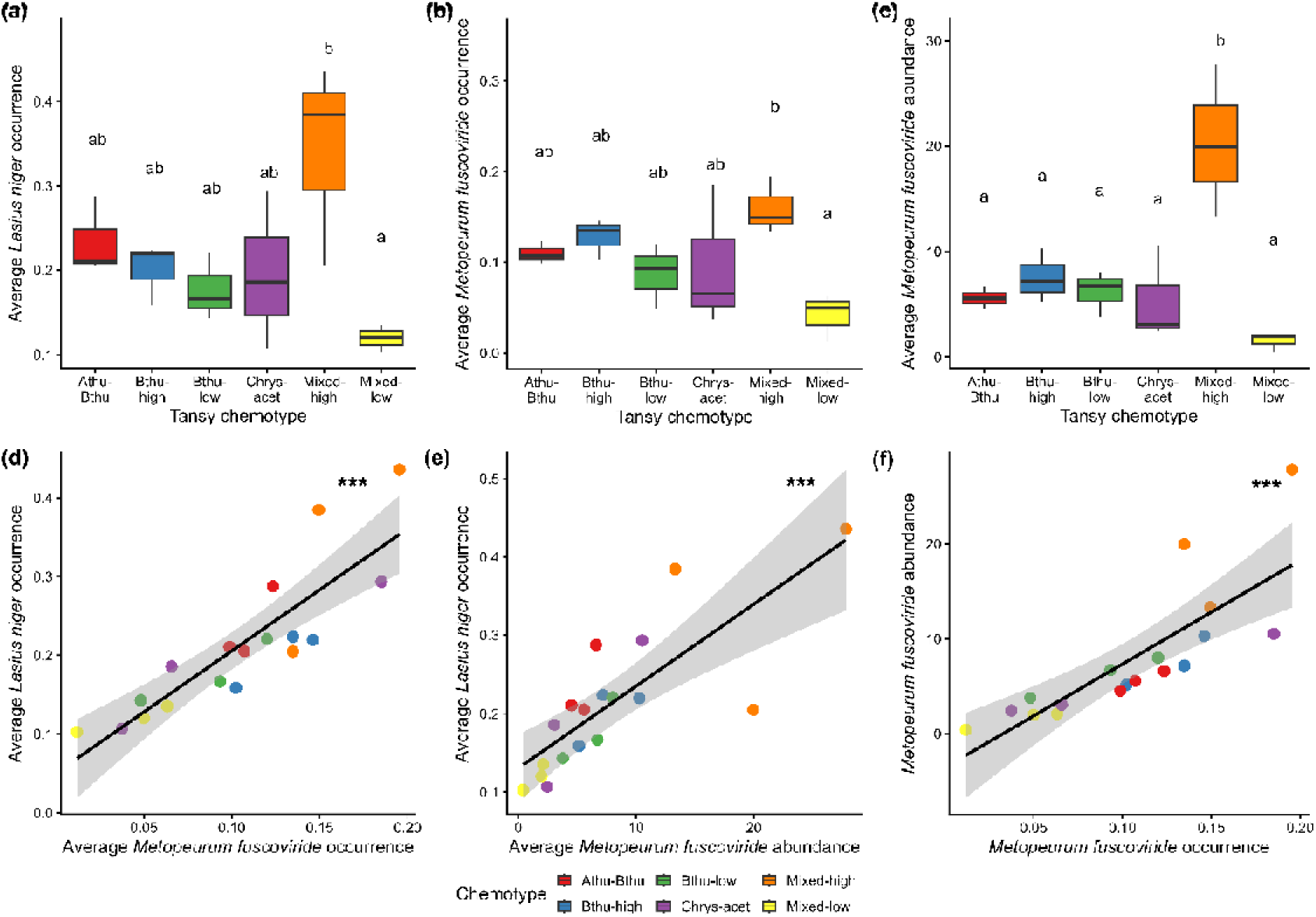
Effect of chemotype on average (a) *Lasius niger* occurrence, (b) *Metopeurum fuscoviride* occurrence and (c) *M. fuscoviride* abundance on Tansy plants within the field. Alphabets represent significant (P<0.05) differences between chemotypes after Tukey’s post hoc analysis. Scatterplots show the relationship between (d) average *M. fuscoviride* occurrence and average *L. niger* occurrence, (e) average *M. fuscoviride* abundance and average *L. niger* occurrence, and (f) average *M. fuscoviride* occurrence and average *M. fuscoviride* abundance. Significance levels are presented as: + = 0.10 > *P* > 0.05, * P<0.05, ** P<0.01, and *** P<0.001. Each daughter (three daughter genotypes for each chemotype) was cloned via cuttings and present 27 – 31 times across different plot conformations. Each dot in Fig. 5 d – f represents the mean value for each daughter across all clones within the field experiment.

## Discussion

Mutualistic relationships in nature do not solely depend on the mutualistic partners involved in them but also on their surrounding environment. In ant-aphid mutualisms, this can take several forms. For instance, the host plant may exert bottom-up pressure that can affect abundance of aphids and their associated organisms, which may influence the ease of ants locating them (Mendes & Cornelissen, 2017; Moreira et al., 2012). However, aphids may favour plants with specific chemical profiles, and ants could eavesdrop on plant-associated cues that reliably yield higher probabilities and abundance of aphid partners (Bristow, 1991). We investigated the role of intraspecific chemodiversity at the plot- and plant-level in influencing the black garden ant, *Lasius niger,* its nesting, patrolling and recruitment behaviour and its commonly observed mutualism with the Pink Tansy aphid, *Metopeurum fuscoviride* on Tansy (*Tanacetum vulgare*) in an established field experiment. Several components of *L. niger* behaviour were subtly but positively influenced by plot-level chemodiversity. The presence of specific individual chemotypes in a plot also had minimal effects on *L. niger* behaviour, including impacts on the location of nests. Importantly, these plot-level chemodiversity impacts on ants varied across the season, with the effects mostly observed in the earlier samplings and disappearing in the later samplings. Additionally, we show that although both the aphids and the ants show preference for specific plant chemotypes, their presence and abundance on plants may be driven most strongly by the presence and abundance of their mutualist partners.

### Distinct aspects of plot-level chemodiversity drive ant behaviour

Before the field experiment was established in May 2021, the site was tilled and vegetation removed, largely homogenizing ant presence through nest disturbance. It is therefore plausible that the observed patterns reflect queen preference while establishing new nests in the summer(s) that followed since. Our studies revealed that among plants of the same species, plot-level chemotype richness has a marginally significant, yet positive impact on the location of ant nests. The choice for nest location is a crucial decision for queen and overall ant colony fitness, which can influence colony access to resources, competition from other colonies, among others (Blüthgen & Feldhaar, 2009; Halder & Annagiri, 2024). Several studies have shown that foundress queens can use plant volatiles to identify suitable plant hosts (Jürgens et al., 2006; Torres & Sanchez, 2017; Razo-Belman et al., 2018). We speculate that the complex blend of volatiles in plots with high chemotype richness may serve as cues for increased host quality and even possible nectar rewards, causing *L. niger* queens to situate their nests within or in proximity to such plots.

Additionally, we found more ant nests near plots having higher stem numbers. During field establishment, all plants had 1 – 3 stems, which increased over time (Ojeda-Prieto, Medina-van Berkum, et al., 2025), and may have influenced queen preference as well, as this generally takes place in high summer, which would have been months after establishment. Tansy stems are inhabited and fed on by mutualistic aphids like *Metopeurum fuscoviride* and *Brachycaudus cardui* which are attended by *L. niger* colonies for honeydew. Therefore, plots with more stems are likely to support a higher abundance of these mutualist aphids, making them suitable sites for placing nests due to resource abundance.

Plot-level chemotype richness did not affect other aspects of ant behaviour like ant patrolling, recruitment time and recruitment numbers. However, we observed variations in ant behaviour across the season (May-July). In late spring (May), when *L. niger* colonies were recovering from hibernation, ant activity was higher in plots in proximity to more nests compared to plots in proximity to less nests, a trend that was not visible in our recruitment assay in later months. This highlights the importance exploring the interactions between Month and the various chemodiversity components on ant behaviour. Although most plots showed relatively rapid recruitment throughout the experiment, we observed variations in recruitment rate at different daily temperatures, which confirms that ant behaviour is regulated by temperature (Cerdá et al., 1998; Markin, 1970; Trigos-Peral et al., 2024). Interestingly, while ants are clearly more active in the warmer months, we do see that within the day, ants are typically most active at the cooler times of day than the warmer times of day, suggesting that they avoid the mid-day heat.

Most of the Tansy chemodiversity metrics had no effect on ant behaviour with the exception of terpenoid abundance. Terpenoid abundance, which is a proxy for terpenoid concentration and emission within plots, had a positive relationship with the number of ants patrolling within plots in May but not in later months. It is possible that *L. niger* uses terpenoid abundance as a positive cue for foraging when the colony is recovering from hibernation in the early stages of its annual life cycle. The main effect of terpenoid abundance on ant activity fades in subsequent months. What drives these patterns is unclear, but plausible explanations could be that in spring, when plants freshly emerge, variation in terpenoid concentration will be most detectable, whereas in summer, when plants have amassed large amounts of leaf biomass, the variation may drown in the vast amounts of terpenoids that are potentially released. Further research is required to understand the spatial behaviour of terpenoids over the course of the growth season, to better understand temporal variation in interaction patterns and how they are mediated by specialized metabolites.

### Presence of specific chemotypes in a plot impacts ant behaviour

We observed significant differences in ant behaviour in the presence of specific Tansy chemotypes. More ant nests were recorded within and in the vicinity of plots having plants of the Chrys-acet chemotype compared to plots without the chemotype. Although chrysanthenyl acetate, the dominant terpenoid in the Chrys-acet chemotype, is not a known attractant of ants or aphids, Ojeda-Prieto, Moreno, et al., (2025) observed a high abundance of *B. cardui* on these plants in the same field experiment in the year of establishment. This aphid is another generalist mutualist of *L. niger,* which often colonizes Tansy plants in early stages, before being gradually replaced by specialist aphids later in the season. It is possible that *L. niger* queens established their nests within and in proximity to plots having the Chrys-acet chemotype in the early years of field establishment, due to the presence of readily available food (honeydew) from, for instance, *B. caudui* and those nests have persisted over the years. This is supported by Blüthgen & Feldhaar, (2009) which states that ants are more likely to establish permanent nests when food is readily available and may be more prone to relocating nests when food is hard to find.

We also found weak effects of the Bthu-low chemotype on ant nesting and patrolling. Bthu-low presence in plots resulted in weak but positive effects on nest numbers. We speculate that this is due to the morphology of plants of this chemotype in the early years of field establishment. In 2021 and 2022, plants of the Bthu-low chemotype were observed to have distinctly tall and thick stems as well as high aboveground biomass (Ojeda-Prieto, Medina - van Berkum, et al., 2025). The same chemotype also had the highest number of flower heads on plants of this chemotype in 2021 (Ojeda-Prieto, Medina-van Berkum et al., 2025). These morphological characteristics could indicate the presence of food resources, potentially attracting founding queens to plots with this chemotype, but this is a hypothesis that remains to be experimentally tested in the Tansy model system. Ant patrolling, however, was negatively affected by the presence of this chemotype, β-thujone the main constituent of this chemotype is a highly toxic feeding deterrent that protect plants from herbivory damage (Wróblewska-Kurdyk et al., 2019; Xie et al., 2019). This toxicity might have made these plots less occupied by mutualist aphids and thereby less attractive to *L. niger* workers for foraging especially when the colony is still recovering and foraging needs to be more precise. Future experimental work is required to test the potential influence of Tansy chemotypes that vary in their chemistry on the nest site selection by ant queens.

### Time of sampling interacts with other drivers of ant behaviour

Ant behaviour was variable in different months, this is evident in how different chemodiversity components impacted ant behaviour across the season. For instance, the influence of nest proximity on number of ants was high in our patrolling assay in May but disappeared in our recruitment assays in subsequent months. We theorise that in late spring (May) when *L. niger* colonies are still recovering from hibernation, they focus on conserving energy and rebuilding the colony. For instance, ants rely on fat reserves accumulated the previous growth season when overwintering. (Enriquez & Visser, 2023). Consequently, these resources are often depleted by the start of a new season. Patrolling and foraging closer to nests may allow surviving workers to conserve the little energy remaining whilst reactivating. Also, ants are ectotherms. As such, they rely on external temperatures for body temperature regulation (Youngsteadt et al., 2023). In late spring, temperatures are still low, particularly in the mornings and evenings, patrolling closer to their nests allows ants to easily return to the safety of their nests if conditions worsen. In subsequent months when atmospheric temperatures get progressively warmer, ants reproduce more rapidly and may forage further from their nests, which may lead to a crowding of the field with workers from many different nests and from further distances.

We found variable responses of ant recruitment to different chemotypes in different months. Plant terpenoid composition differs at different developmental stages (Ziaja & Müller, 2025; Flamini et al., 2013), the variations in terpenoid blends emitted by the different Tansy chemotypes at their different growth stages may have resulted in different responses of ants and aphids to Tansy chemotypes across the season. These differences were more pronounced on time to first recruitment to bait because of the additional impact of temperature. Environmental factors like temperature can influence terpenoid emission by plants and their perception by insects (Malik et al., 2023; Ruano et al., 2000; Yuan et al., 2009).

### Plant chemotypes differ in ant occurrence, mediated via aphid occurrence and aphid abundance

When we investigated ant and aphid presence and abundance at the level of individual plants, we observed the highest incidence of both ants and aphids on plants of the Mixed-high chemotype, suggesting a clear preference for this chemotype. The disparity in *L. niger* responses to chemodiversity between plots and plants, as well as the strong relationship with aphid patterns on plants, suggests that ant occurrence on plants may be more dependent on the presence and abundance of their mutualist *M. fuscoviride* than on plant and plot chemistry. Notably, we also found the lowest incidence of ants and aphids on the Mixed-low chemotype. Although both Mixed-high and Mixed-low are heterogeneous chemotypes characterised by high terpenoid evenness and diversity, differences exist in their terpenoid profiles and concentrations. Overall terpenoid concentrations are lower in the Mixed-low chemotype compared to Mixed-high chemotype. Moreover, closer examination of their profiles also reveals some terpenoids that are not shared between the two chemotypes. These factors may have resulted in the different responses of ants and aphids to these chemotypes. Mixed-low chemotypes may be low quality hosts of aphids. According to (Neuhaus-Harr et al., 2025), more winged individuals of a different specialist aphid, the Tansy aphid, *M. tanacetaria,* were found on the Mixed-low chemotype. Production of winged morphs by aphids can be induced in response to environmental stressors like reduced host plant quality (Mehrparvar et al., 2013; Johnson, 1965). It is likely that this low host quality may have led to aphids avoiding this chemotype, this low aphid numbers will eventually lead to reduced number of tending ants. Conversely, Neuhaus-Harr et al., (2025) also found the highest colony growth of aphid *M. tanaceteria* on the Mixed-low chemotype, which suggests that different aphid species may have varying responses to the same terpenoid profile.

## Conclusion

In summary, our findings provide further insights into the effect of plant and plot chemistry on ant behaviour and ant-aphid mutualisms. We show that although plot-level chemotype richness has marginally significant effects on *L. niger* nesting and does not affect most components of *L. niger* worker behaviour. Individual plant chemotypes and other inherent chemical diversity elements, however, clearly influence ant behaviour variably across the season. Our findings suggest that, more importantly, *L. niger* incidence on plants may depend more on *M. fuscoviride* presence than on plant chemistry.

## Supporting information

Supplements to Setordjie et al.

## Acknowledgements

We are grateful for support of the Jena Experiment management, in particular Anne Ebeling and Nico Eisenhauer for allowing us to maintain our experiment on-site, and to the local gardening team for their excellent support and allowing us to use their equipment where needed.

## Author contributions

W.W. Weisser designed the experiment. R. Heinen propagated the plants and R. Heinen, L. Ojeda-Prieto and W. W. Weisser set-up the field. E. A. Setordjie and L. Ojeda-Prieto conducted surveys and maintained the field experiment. E. A. Setordjie organized and analysed field data. L. Ojeda-Prieto analysed leaf terpenoid metrics. E. A. Setordjie wrote the manuscript with guidance the of R. Heinen. L. Ojeda-Prieto, and W.W. Weisser provided critical feedback on earlier drafts. All authors contributed to the final version and approved the final submission.

## Conflict of interest

The authors declare no conflict of interest to disclose.

Link to Biorender license: https://biorender.com/off80or

