## Supplements to Setordjie et al. for "Aphid presence and abundance, more than variation in leaf terpenoid profiles at the plant and plot-level, drive ant behaviour on the perennial forb *Tanacetum vulgare*"

**Supplementary tables**

**Table S1:** Model output for the effect of plot-level chemotype richness.

Degrees of freedom (df), Wald’s Chi-square statistics and p-values are reported in the table.

NMP = Number of nests within a meter of plot, CR = Plot-level chemotype richness, NA = not applicable

|  |  | NMP | Patrolling | Recruitment time (bait) | Total number of ants (bait) |
| --- | --- | --- | --- | --- | --- |
|  | d.f. | X^2^ (p-value) | X^2^ (p-value) | X^2^ (p-value) | X^2^ (p-value) |
| NMP | 1 | NA | 7.2 (0.007) | 1.1 (0.293) | 0.0 (0.831) |
| Temperature | 1 | NA | NA | 31.3 (<0.001) | 0.0 (0.914) |
| Number of stems | 1 | 4.6 (0.032) | 0.8 (0.365) | 0.7 (0.393) | 1.4 (0.239) |
| CR | 1 | 3.6 (0.058) | 0.0 (0.962) | 0.0 (0.912) | 0.2 (0.623) |
| Month | 2 | NA | NA | 347.8 (< 0.001) | 31.1 (< 0.001) |
| CR*Month | 2 | NA | NA | 0.3 (0.849) | 1.1 (0.583) |

**Table S2**: Model output for the effects of plot-level chemodiversity metrics

Degrees of freedom (df), Wald’s Chi-square statistics and p-values are reported in the table. NMP = Number of nests within a meter of plot, NA = not applicable

|  |  | NMP | Patrolling | Recruitment time (bait) | Total number of ants(bait) |
| --- | --- | --- | --- | --- | --- |
|  | d.f. | X^2^ (p-value) | X^2^ (p-value) | X^2^ (p-value) | X^2^ (p-value) |
| NMP | 1 | NA | 6.0 (0.014) | 0.9 (0.349) | 0.1 (0.706) |
| Temperature | 1 | NA | NA | 33.3 (<0.001) | 0.0 (0.967) |
| Number of stems | 1 | 4.5 (0.033) | 0.5 (0.465) | 0.0 (0.828) | 0.4 (0.519) |
| Terpenoid richness | 1 | 0.5 (0.482) | 0.4 (0.510) | 0.1 (0.746) | 2.1 (0.143) |
| Terpenoid Shannon diversity | 1 | - | - | - | - |
| Terpenoid Hill Evenness | 1 | 0.1 (0.713) | 0.0 (0.983) | 0.6 (0.445) | 1.6 (0.212) |
| Total terpenoid abundance | 1 | 0.9 (0.350) | 10.7 (0.001) | 0.7 (0.402) | 0.1 (0.704) |
| Month | 2 | NA | NA | 412.7 (<0.001) | 42.7 (<0.001) |
| Terpenoid richness * Month | 1 | NA | NA | 3.3 (0.188) | 2.1 (0.331) |
| Terpenoid Shannon diversity * Month | - | - | - | - | - |
| Terpenoid Hill Evenness * Month | 1 | NA | NA | 10.9 (0.004) | 0.6 (0.735) |
| Total terpenoid abundance * Month | 1 | NA | NA | 50.9 (<0.001) | 7.3 (0.026) |

**Table S3**: Model output for the effects of chemotype presence.

Degrees of freedom (df), Wald’s Chi-square statistics and p-values are reported in the table. NMP = Number of nests within a meter of plot, NA = not applicable

|  |  |  | NMP | Patrolling | Recruitment time (bait) | Total number of ants (bait) |
| --- | --- | --- | --- | --- | --- | --- |
|  |  | d.f. | X^2^ (p-value) | X^2^ (p-value) | X^2^ (p-value) | X^2^ (p-value) |
| Athu-Bthu | NMP | 1 | NA | 7.1 (0.008) | 1.1 (0.298) | 0.0 (0.826) |
|  | Temperature | 1 | NA | NA | 31.3 (<0.001) | 0.0 (0.923) |
|  | Number of stems | 1 | 4.6 (0.031) | 0.7 (0.396) | 0.7 (0.401) | 1.2 (0.265) |
|  | Chemotype presence | 1 | 1.1 (0.295) | 0.3 (0.601) | 0.0 (0.967) | 0.8 (0.364) |
|  | Month | 1 | NA | NA | 415.8 (<0.001) | 42.1 (<0.001) |
|  | Chemotype presence * Month | 1 | NA | NA | 2.4 (0.297) | 1.6 (0.453) |
| Bthu-high | NMP | 1 | NA | 7.6 (0.006) | 1.1 (0.298) | 0.2 (0.693) |
|  | Temperature | 1 | NA | NA | 31.4 <0.001) | 0.0 (0.937) |
|  | Number of stems | 1 | 4.2 (0.040) | 0.8 (0.368) | 0.7 (0.396) | 1.5 (0.226) |
|  | Chemotype presence | 1 | 0.1 (0.722) | 0.2 (0.634) | 0.0 (0.866) | 2.0 (0.159) |
|  | Month | 1 | NA | NA | 416.2 (<0.001) | 42.4 (<0.001) |
|  | Chemotype presence * Month | 1 | NA | NA | 3.2 (0.197) | 4.9 (0.088) |
| Bthu-low | NMP | 1 | NA | 9.3 (0.002) | 1.1 (0.299) | 0.0 (0.903) |
|  | Temperature | 1 | NA | NA | 31.1 (<0.001) | 0.0 (0.908) |
|  | Number of stems | 1 | 5.2 (0.022) | 1.3 (0.261) | 0.7 (0.398) | 1.2 (0.283) |
|  | Chemotype presence | 1 | 2.7 (0.099) | 2.7 (0.099) | 0.0 (0.965) | 1.0 (0.308) |
|  | Month | 1 | NA | NA | 418.7 (<0.001) | 42.1 (<0.001) |
|  | Chemotype presence * Month | 1 | NA | NA | 47.5 (<0.001) | 1.3 (0.533) |
| Chrys-acet | NMP | 1 | NA | 5.7 (0.017) | 0.5 (0.460) | 0.0 (0.906) |
|  | Temperature | 1 | NA | NA | 31.4 (<0.001) | 0.0 (0.924) |
|  | Number of stems | 1 | 5.4 (0.020) | 0.5 (0.489) | 0.4 (0.518) | 1.1 (0.288) |
|  | Chemotype presence | 1 | 4.0 (0.045) | 2.2 (0141) | 1.5 (0.224) | 0.7 (0.387) |
|  | Month | 1 | NA | NA | 419.0 (<0.001) | 42.5 (<0.001) |
|  | Chemotype presence * Month | 1 | NA | NA | 98.7 (<0.001) | 5.8 (0.054) |
| Mixed high | NMP | 1 | NA | 7.6 (0.006) | 1.2 (0.282) | 0.1 (0.757) |
|  | Temperature | 1 | NA | NA | 31.4 (<0.001) | 0.0 (0.919) |
|  | Number of stems | 1 | 4.1 (0.043) | 0.4 (0.534) | 1.1 (0.284) | 1.0 (0.324) |
|  | Chemotype presence | 1 | 0.0 (0.952) | 1.0 (0.308) | 0.9 (0.342) | 0.5 (0.495) |
|  | Month | 1 | NA | NA | 416.0 (<0.001) | 42.2 (<0.001) |
|  | Chemotype presence * Month | 1 | NA | NA | 5.2 (0.074) | 2.5 (0.293) |
| Mixed low | NMP | 1 | NA | 8.0 (0.005) | 1.2 (0.274) | 0.1 (0.727) |
|  | Temperature | 1 | NA | NA | 31.5 (<0.001) | 0.0 (0.928) |
|  | Number of stems | 1 | 2.4 (0.119) | 0.1 (0.764) | 0.1 (0.732) | 0.7 (0.418) |
|  | Chemotype presence | 1 | 0.4 (0.540) | 1.4 (0.242) | 0.8 (0.357) | 0.4 (0.550) |
|  | Month | 1 | NA | NA | 414.3 (<0.001) | 42.1 (<0.001) |
|  | Chemotype presence * Month | 1 | NA | NA | 14.7(<0.001) | 1.9 (0.384) |

**Table S4**: Summary of simple linear regression models (LM) assessing the effect of Tansy plant chemotype on Average ant occurrence, Average aphid occurrence and Average aphid abundance Degrees of freedom (d.f.), F statistics and p-values are reported in the table. ^#^Effect of chemotype on response variables when only plants with aphid presence are assessed. NA = not applicable.

|  | d.f. | Ant occurrence | Aphid occurrence | Aphid abundance |
| --- | --- | --- | --- | --- |
|  |  | F (p-value) | F (p-value) | F (p-value) |
| Chemotype | 1 | 3.45 (0.037) | 2.903 (0.061) | 8.83 (0.001) |
| Chemotype^#^ | 1 | 1.26 (0.34) | NA | 13.69 (<0.001) |

**Supplementary figures**

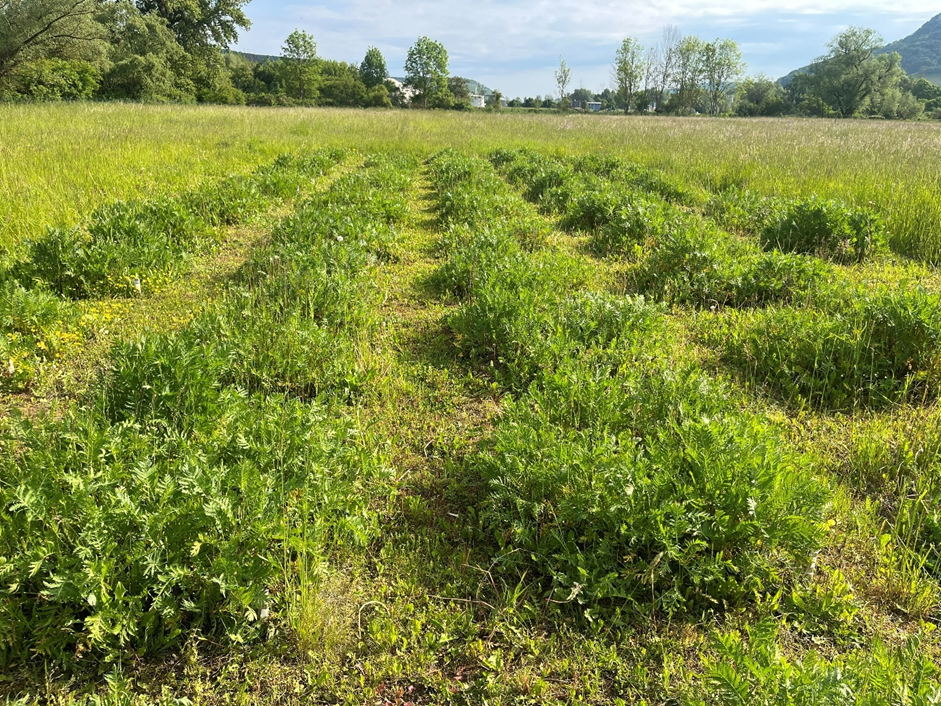

**Figure S1:** Field layout in the Jena experiment

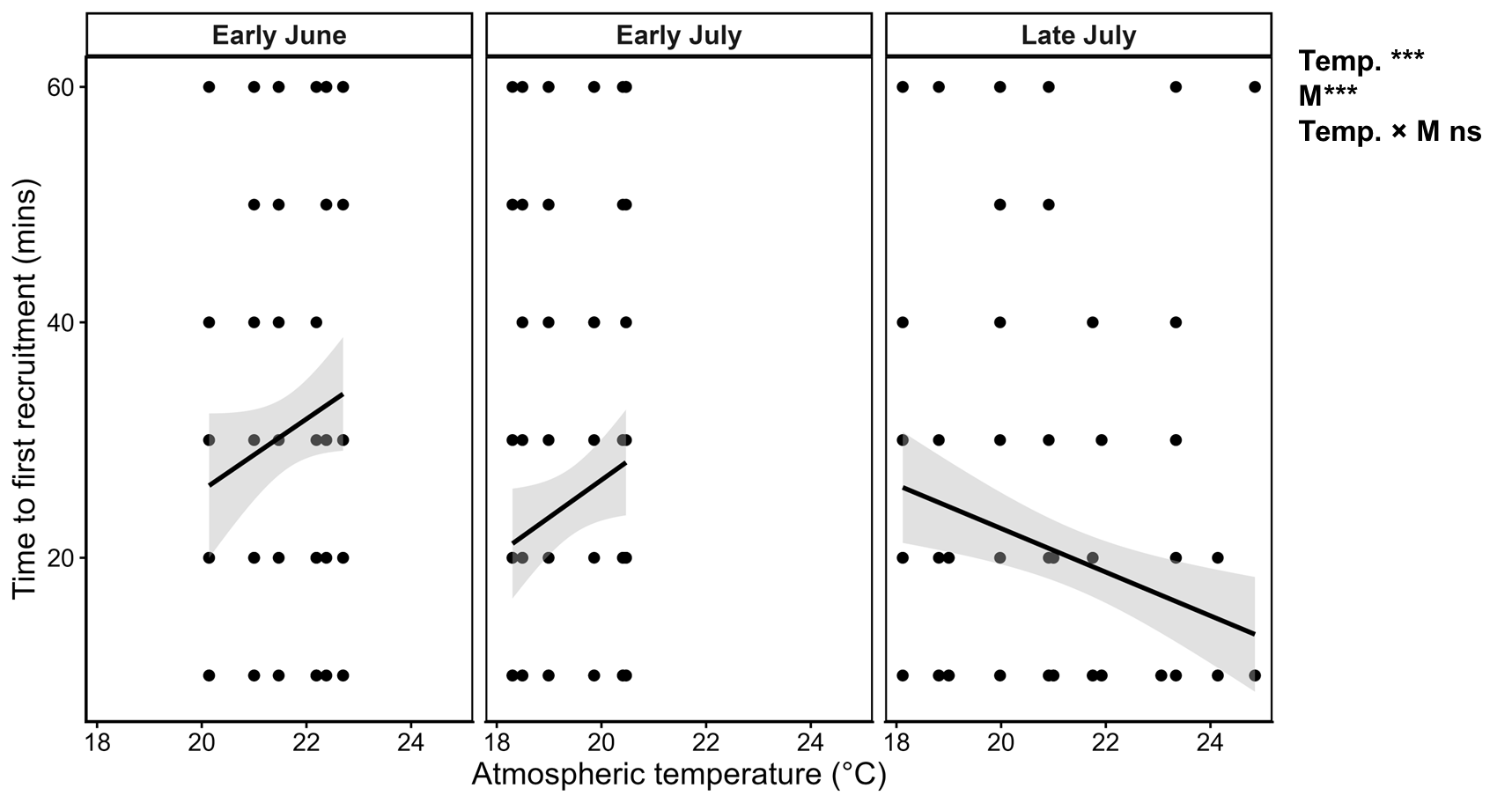

**Figure S2**: Effect of Atmospheric temperature (Temp), month of observation (M) and their interaction on time to first ant recruitment throughout the course of sampling. Significance levels are presented as: P<0.1, * P<0.05, ** P<0.01, and *** P<0.001 on the top right corner of figure.

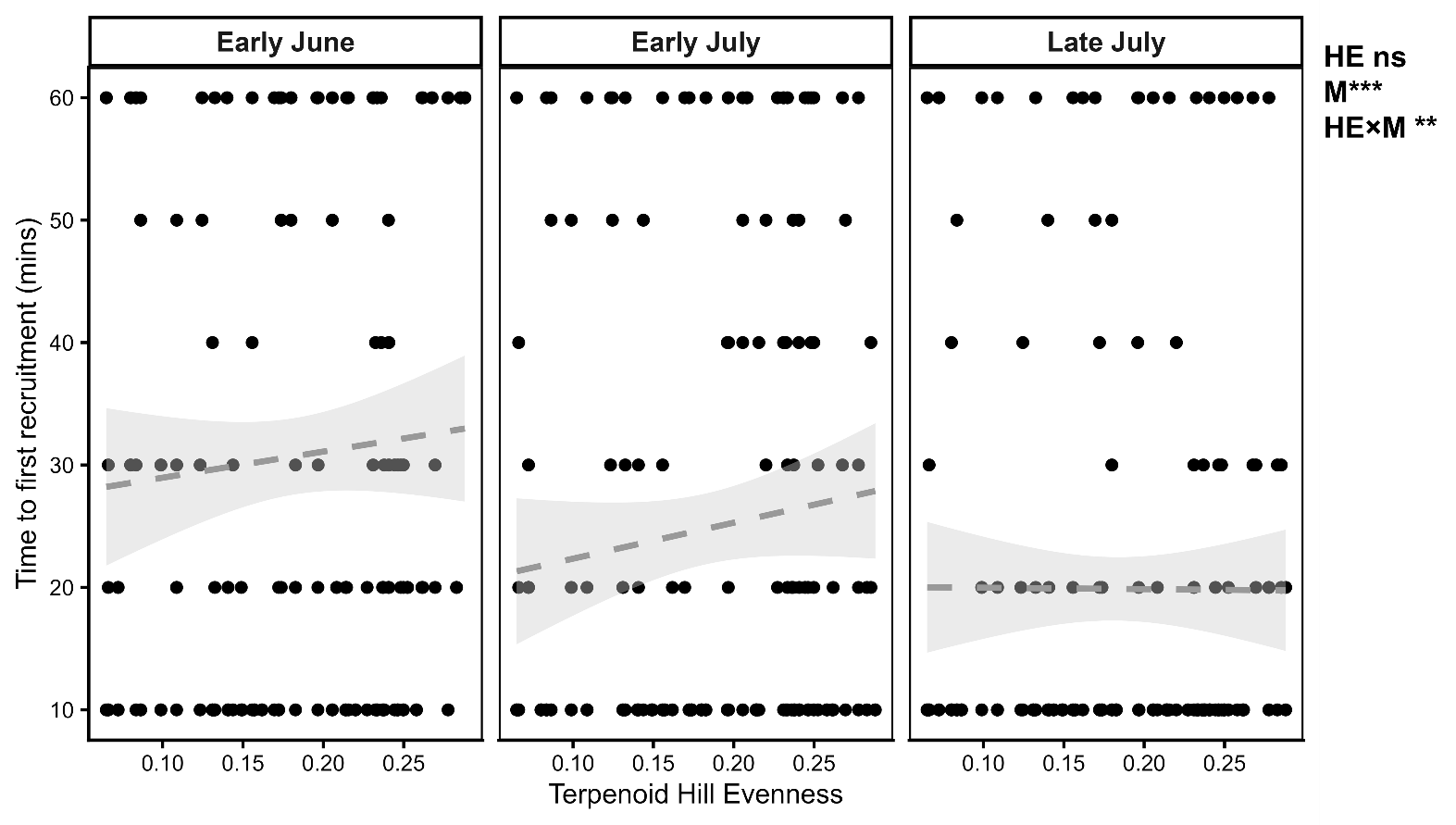

**Figure S3:** Effect of Terpenoid Hill evenness (HE), month of observation (M) and their interaction on time to first ant recruitment on honey baits. Significance levels are presented as: ns = not significant, P<0.1, * P<0.05, ** P<0.01, and *** P<0.001 on the top right corner of figure.

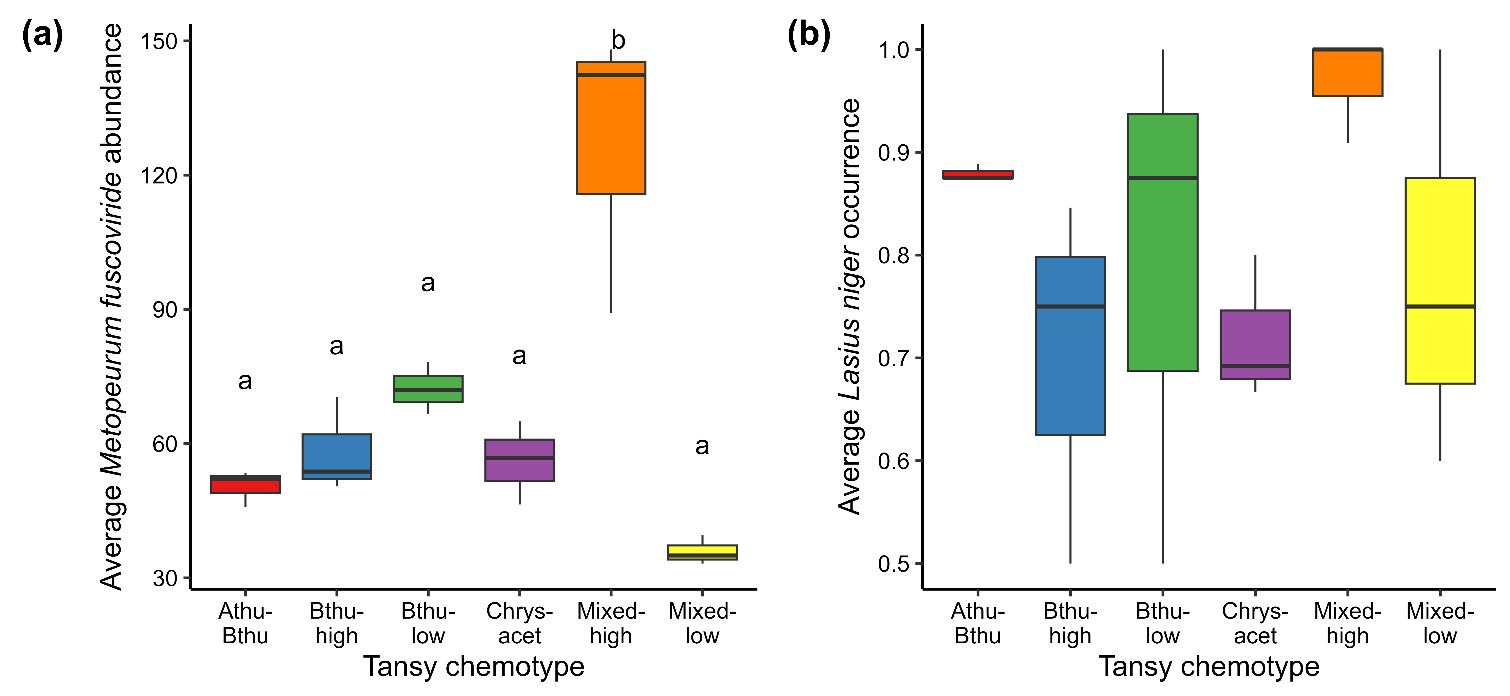

**Figure S4**: Effect of chemotype on average (a) *Lasius niger* occurrence (b) *Metopeurum fuscoviride* occurrence and (c) *Metopeurum fuscoviride* abundance including only those Tansy plants that were colonized and thus had aphid presence within field. Alphabets represent significant (p<0.05) differences between chemotypes after Tukey’s post hoc analysis.
